# A pilot study on the biological applications of indole alkaloids derived from Nagaland biodiversity

**DOI:** 10.1101/2025.01.08.632032

**Authors:** Aben Ovung, A. Mavani, Gourisankar Roymahapatra, Jhimli Bhattacharyya

## Abstract

The north-eastern Indian state, Nagaland with its rich biodiversity, is well known for traditional, natural medicines. However, the scientific basis of the working principle(s) of such drugs are yet to be explored. Plant alkaloids, the nitrogen-containing organic molecules, are traditionally been used for various medicinal purposes from ages. There is a need for drug discovery from natural sources to create a broader drug portfolio for human use. And, Nagaland medicinal plant based bioactive molecules have huge potential for the same. As a pilot study, this work aims to study the *in silico* absorption, distribution, metabolism and excretion (ADME) properties, toxicity prediction and structural properties of the indole alkaloid molecules, Ajmalicine (AJM) and Serpentine (SER) and its docking interaction pattern with various forms of DNA. ADME analysis confirms that the alkaloids meet the criteria of a standard drug. The ProTox II prediction showed the probability of the toxicity profile of the input compounds. Density functional theory (DFT) revealed the HOMO-LUMO energy gap of the drug molecules along with the quantum chemical parameters from the optimized geometry of the ligands. Molecular docking (MD) analysis showed successful binding of the molecules into the groove region of the various forms of DNA structure for both the cases. The results of this theoretical pilot study provides information that will be helpful in the development of effective and modified drugs for medicinal and pharmaceutical purposes as well as in the planning of practical laboratory experiments.

## 1. Introduction

Nagaland, an integral part of North-eastern India, is a biodiversity hotspot in the country.[1] The state is home to several ethnic tribes that still use various conventional practices, such as traditional medicinal practices in which plants play an important role. The ethnic communities and their traditional knowledge on the use of herbal medicines have survived over the centuries. The rich biodiversity of Nagaland ensures enormous potential of various medicinal and allied plants growing freely in nature.[2] The direct use of traditional medicines of plant origin, such as leaves, roots, bark, etc., is widespread, but the exact characterization and scientific background of these natural medicines, as well as their mechanism of action, still need to be explored. Many plants used in traditional medicine of Nagaland are found to contain indole alkaloids as one of the active component.[3][4] It is learned that, alkaloids have important physiological effects and significant therapeutic properties in humans and animals.[5, 6][7] Alkaloids are a large class of naturally occurring organic molecules that contain one or more nitrogen atoms in their structure.[8, 9] Alkaloids play a very important role in pharmacological effects on human health, such as anticancer, antimalarial, antihypertensive, antidiabetic effects etc.[10] Alkaloids can have a direct effect on the central nervous system and also influences nucleic acid, membrane permeability and proteins.[11]

Medicines derived from plants represent an important source of innovation for novel active ingredients that greatly benefit human health and well-being.[12] Plants consist of a variety of components that can be used for the production and development of various drug molecules. For the present study, we have tried to focus on the indole alkaloids; Ajmalicine (AJM) and Serpentine (SER) which are actively found in the plants like *Catharanthus roseus* (common name: Periwinkle; Nagaland local name in Ao Changi language: Tsǖinri naro)*, Rauvolfia serpentina* (common name: Indian Snakeroot/Sarpagandha; local name: Per-mozutong). Figure 1 (A) and (B) show the structure of AJM and SER which are known as antihypertensive drug and are commonly used for the treatment of high blood pressure,[13–15] loss of memory, cognitive deficit and hypertension.[16]

**Figure 1.**
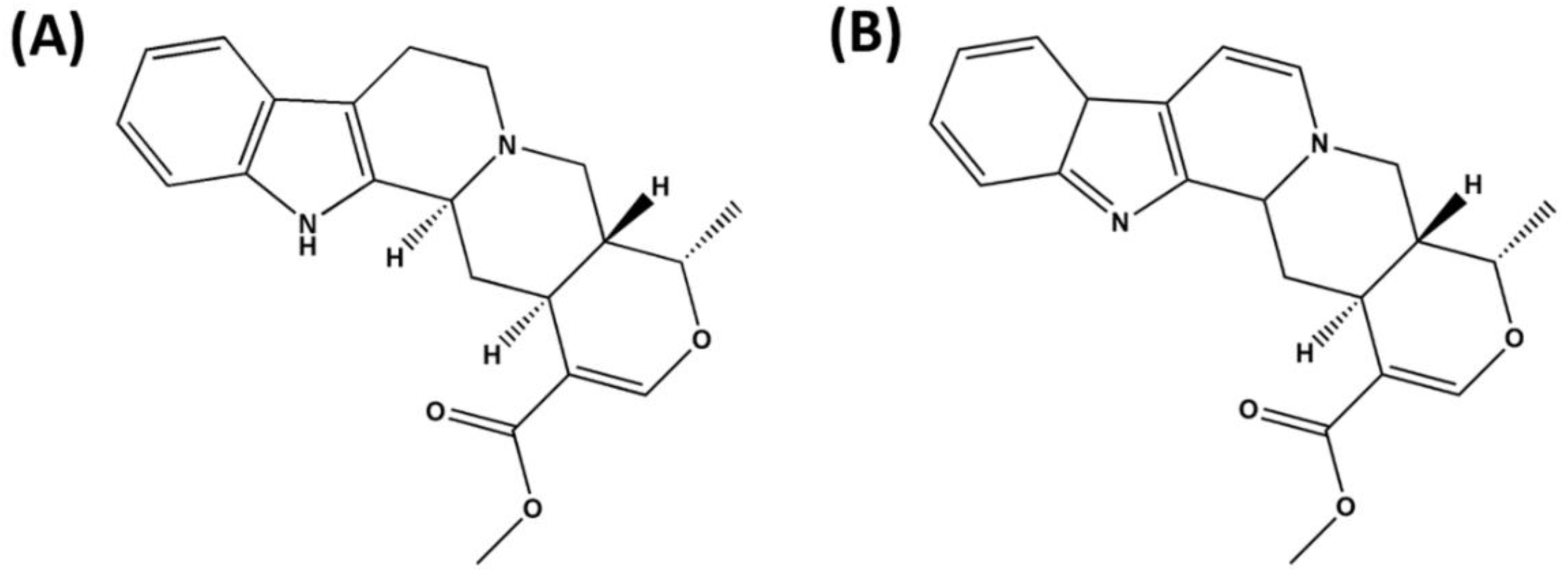
Chemical structures of (A) Ajmalicine (AJM), (19α)-16, 17-didehydro-19-methyloxayohimban-16-carboxylic acid methyl ester and (B) Serpentine (SER), (19α)-16-(Methoxycarbonyl) - 19 - methyl - 3, 4, 5, 6, 16, 17-hexadehydro-18-oxayohimban-4-ium-1-ide.

*In silico* studies using various bioinformatics programs or web servers are currently the primary method for reducing the time required for drug discovery and development. Computational strategies are of great value for the identification and development of new promising compounds.[17, 18] Wide range of small molecules/ligands interfere the biological effects by interacting reversibly with nucleic acid [19][20]. Deoxyribonucleic acid (DNA) is a most frequently targeted bimolecular system by therapeutic agents [21]. Therefore, the study of drug-DNA interactions is an active area of research that is driving the development of new modified, promising, and effective drugs for clinical use [22]. The interaction pattern can be either classical, non-classical, furrow or electrostatic [23, 24]. Studies on molecular modeling; theoretical and computational techniques play an important role in understanding the behavior and mechanisms of the structure of molecule and complex formation in a non-covalent manner.[25][26] The results of the theoretical study provide protocols to complement experimental data for structural activity correlation in drug-DNA complexes.[27] Various studies on the extraction and use of AJM and SER have been investigated by many researchers [16][28][29].

This paper presents a detailed *in-silico* study of the two biologically important alkaloid components (AJM and SER) using SwissADME, ProTox-II, Density Functional Theory (DFT) and its binding/interaction pattern with various forms of DNA (such as duplex, hairpin duplex, triplex and quadruplex DNAs). The study helps us to understand the physical and chemical properties of the two studied indole alkaloid molecules and to predict/identify the best target region for DNA. Overall, this study provides a fundamental knowledge, properties and molecular mechanism of indole alkaloid molecules (AJM and SER) and their interaction patterns with different forms of DNA. The study may further help in the discovery of potential small molecules present in the northeast region of India and their therapeutic effect in terms of interaction with bio-macromolecules.

## 2. Computational details

### 2.1. SwissADME *in-silico* study

SwissADME software (http://www.swissadme.ch/) is a free online platform for determining absorption, distribution, metabolism and excretion (ADME) properties. Parameters like Physicochemical, water solubility, lipophilicity, pharmacokinetics, drug-likeness and medicinal chemistry of small molecule for drug discovery and development were investigated through the study.[30–32] The chemical structure of the compounds, AJM (https://pubchem.ncbi.nlm.nih.gov/compound/Ajmalicine) and SER (https://pubchem.ncbi.nlm.nih.gov/compound/Serpentine-_alkaloid) were downloaded from PubChem data bank in SDF format. The files were then imported to SwissADME web page and converted into molecular sketcher based on ChemAxon’s Marvin JS followed by ADME calculation using default parameters.

### 2.2. *In-silico* prediction of toxicity

The toxicities of the alkaloid drugs, AJM and SER were predicted using ProTox-II, a freely available *in-silico* toxicity prediction web server (https://tox-new.charite.de/protox_II/index.php?site=compound_input). The chemical structures were developed using the smiles into the ProTox-II web server for the analysis. The prediction involved are of the machine learning algorithm based models. The platform is classified into four different groups namely, organ toxicity (one model), toxicity endpoints (four models), Tox21 Nuclear Receptor Signaling Pathways (seven models) and Tox21 Stress Response Pathways (four models).

### 2.2. Density Functional Theory (DFT) Study

DFT calculations were adopted to model the electronic structures and transitions of the alkaloid drugs, AJM and SER. GAUSSIAN 09W program package was used by considering the most probable structure for the calculations utilizing the hybrid DFT functional B3LYP function and keeping the basis sets as 6-31G (d, p).[33, 34] The highest occupied molecular orbital (HOMO) and lowest unoccupied molecular orbital (LUMO) energies of the electronic properties and other energies related to the drug structures were calculated using B3LYP method of the time-dependent DFT (TD-DFT).

### 2.3. Molecular Docking

Docking interaction analyses using Autudock 1.5.6 software were performed to explore the most feasible binding site, mode and affinity of the alkaloid (ajm and ser) drugs with DNA. The structures of the different forms of DNA was extracted from RCSB Protein Data Bank having PDB ID: 1BUT (https://www.rcsb.org/structure/1BUT) for duplex DNA, 4F41 (https://www.rcsb.org/structure/4F41) for hairpin DNA, 136D (https://www.rcsb.org/structure/136D) and 3SC8 (https://www.rcsb.org/structure/3SC8) for quadruplex DNA while the alkaloid drugs (AJM and SER) were obtained from the frequency optimization of the DFT analysis. The Autodock software utilizes Lamarckian Genetic Algorithm (LGA). To prepare, DNA and the ligands for docking, water molecules were removed and polar hydrogen atoms, Gasteiger charges were added. Autodock tools were used to assign the rotatable bonds in the ligands. In order to locate the binding site of the drugs in DNA, they were bordered and set in the grid box to 80, 80, and 80 Ǻ along the x-, y-, and z-axis with a 0.36 Ǻ grid spacing. The resultant minimum energy docked model was chosen for further analysis and viewed in chimera and discovery studio for better visualization.

## 3. Results and discussions

### 3.1. SwissADME In-silico Study

#### 3.1.1. Boiled-Egg and Bioavailability Radar Study

The pharmacokinetic properties of the alkaloid drugs AJM and SER were studied using boiled egg and bioavailability radar model (figure 2). Figure 2A shows the assessment model for boiled egg. The white area represents a high probability of passive gastrointestinal absorption (HIA) in the gastrointestinal tract, while the yellow area represents a high probability of penetration of the blood-brain barrier (BBB).[35] The analysis predicted that both the drug molecules exhibited BBB penetration. BBB is a micro-vascular endothelial cell layer that surrounds the central nervous system (CNS) which prevents the small molecules from entering the brain.[36] The delivery of drugs to CNS, limiting the access to brain parenchyma is a major challenge in the treatment of neurological diseases[37] and these small molecules may be helpful in the discovery and development of neurological therapeutic drugs. The blue shaded color representing the two drug signify the molecules as P-glycoprotein (PGP+) substrate which are predicted as effectively effluxed by P-glycoprotein. P-glycoprotein plays an important role in drug absorption and disposition as a carrier mediated transporter.[38] They are widely distributed throughout the body (small intestine, blood brain barrier capillaries, kidney, liver etc.) and can bind with a variety of substrate. The bioavailability radar in figure 2 (B and C) shows pink colored zone which accounts for six physicochemical properties (lipophilicity, size, polarity, solubility, saturation and flexibility) into consideration. The optimal range includes lipophilicity: XLOGP3 between −0.7 and +5.0, size: MW between 150 and 500 g/mol, polarity: TPSA between 20 and 130 Å^2^, solubility: log *S* not higher than 6, saturation: fraction of carbons in the sp^3^ hybridization not less than 0.25, and flexibility: no more than 9 rotatable bonds.[39] The drugs from the radar indicate that they fall within the permissible range/parameter of a standard drug.

**Figure 2.**
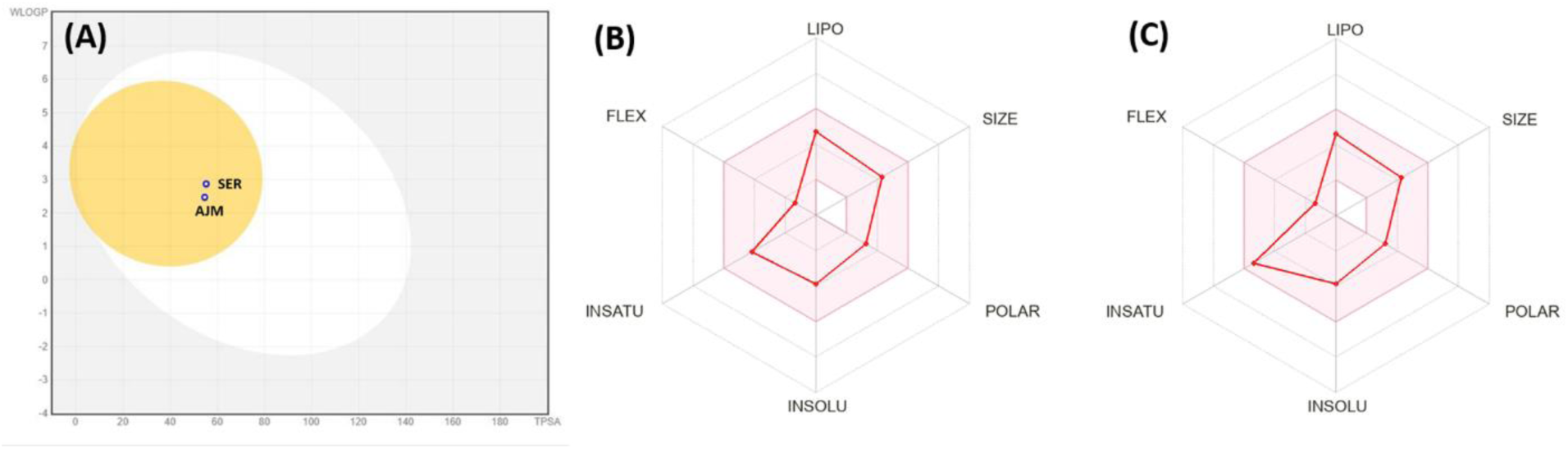
Schematic representation of (A) Boiled-Egg evaluation of passive gastrointestinal absorption (HIA) and Blood Brain Barrier (BBB) penetration (B) and (C) represents Bioavailability Radar for drug likeness of the molecules, AJM and SER, respectively.

#### 3.1.2. ADME properties

Table S1 to S6 represents the ADME properties (physicochemical, lipophilicity, solubility, pharmacokinetics, drug-likeness and medicinal chemistry, respectively) of the alkaloid drugs, AJM and SER. Physicochemical properties like description of number of atoms, hydrogen bonds, molar refractivity (MR) and topological polar surface area (TPSA) are reported in Table S1. The data obtained are useful descriptors for many models to estimate brain access and gastrointestinal absorption.[36, 40] Lipophilicity characteristics of the molecules were determined to evaluate the arithmetic mean of consensus log *P*_o/w_ (Clog P) predicted by five methods (iLOGP, XLOGP3, WLOGP, MLOGP, SILICOS-IT). The logarithm partition coefficient between *n-*octanol/water (log *P*_o/w_) is the measure of lipophilicity.[41] The main action to be considered from the above evaluation is the measure of toxicity in short-term animal studies. The compounds have been categorized into three group based on their condition that includes, highly toxic when Clog P>3 (high lipophilicity) and TPSA<75, non-toxic when Clog P<3 (non lipophilicity) and TPSA>75, neutral when Clog P<3 (non lipophilicity) and TPSA<75.[42, 43] The evaluation suggest that high lipophilicity with low polar surface area has high chances of becoming toxic. The data obtained from the drugs (AJM and SER) reveals that they are neutral molecule with low lipophilicity and polar surface area.

The solubility of drug molecules are divided into four types – high soluble and high permeable, low soluble and high permeable, high soluble and low permeable, low soluble and low permeable.[44] Drug water solubility is predicted by three topological approaches that includes ESOL, Ali and SILICOS-IT models (Solubility class: Log S Scale: Insoluble<-10 poorly<-6, moderately<-4 soluble<-2 very<0<highly). The SILICOS-IT approach represents the model where the linear coefficient is corrected by molecular weight (R^2^ = 0.75).[45] The investigated alkaloids showed soluble for the first two model and moderately soluble for the latter, Table S3.

Metabolization is an important and interesting pharmacokinetics process to predict via *in silico* methods. Cytochrome p450 (CYP) is a family of enzyme that acts actively in two phases of metabolism, Phase I reactions (oxidation, reduction, hydrolysis and O/N de-alkylation) Phase II reactions (Conjugation by glucuronidation, acetylation, sulphation and methylation).[46] CYP enzyme bio-transforms the molecules through five different isoforms (CYP1A2, CYP3A4, CYP2C9, CYP2C19, CYP2D6). The resulting molecules are indicated in Table S4 as “Yes” or “No” if the molecules under investigation are expected to act as substrates for CYP. The reported models were validated and constructed using machine learning technique, Support Vector Machines (SVM). The skin permeation coefficient (Log *K*_p_) is also expressed, resulting in lower skin permeability when the value for a molecule is more negative. After the pharmacokinetic evaluation, the molecules were examined for druglikeness rule and bioavailability score. The analysis of oral bioavailability drugs were based on Lipinski’s, Ghose filter, Vebers’s, Egan’s, and Muegge’s rules. The result of the two alkaloid molecules, AJM and SER showed good and acceptable prediction with a high bioavailability score and no violation for all the cases. Lastly, medicinal chemistry properties was analyzed for the alkaloids to support the drug discovery venture. Pan assay interference compounds (PAINS) considers chemical compounds that shows potent response in assays irrespective of the protein targets which can react with numerous biological targets non-specifically. Brenk account for compounds which are small and less hydrophobic not defined by “Lipinski’s rule of 5” while leadlikeness is the concept of designing lead high affinity optimization exposed to chemical modifications with decrease in size and increase in lipophilicity.[47] The investigated molecules showed 1 alert (Pains) and 1 violation (leadlikeness) for AJM while SER passed all the test of medicinal chemistry properties. The score of synthetic accessibility is related to synthesis ease of the compounds whose score ranges from 10 (very difficult) to 1 (very easy). The synthetic score obtained was 4.32 for SER, little lesser and easier to synthesize as compared to that of AJM (4.57), also suggesting that the molecules are not difficult to synthesize.

### 3.2. *In Silico* Potential Toxicity Study

The *in silico* web server platform ProTox-II enables the prediction of a large number of toxicity endpoints incorporating molecular similarity, pharmacophores, fragment propensities, and machine learning models that provide insights into the potential molecular mechanism.[48–50] The prediction helps to generate new hypotheses to gain insights into the mechanisms of toxicity and initiate follow up experimental studies. Various toxicity end points from the ProTox-II server were predicted that includes acute toxicity, hepatotoxicity, cytotoxicity, carcinogenicity, immunotoxicity, mutagenicity, Tox21 pathways (Table S7). The acute (oral) toxicity of the investigated drugs projected AJM as LD50 (lethal dose) of 300 mg/kg with 68.07% prediction accuracy whereas SER showed LD50 of 198 mg/kg with 54.26%. Both the drugs obtained class III level prescribed as toxic after swallowing (50 < LD50 ≤ 300).[51, 52] Figure 3 represents the toxicity radar plot which provides the assessment of comparison between the average probability score of the active compounds from the training set to that of the input alkaloid compounds.[53] The orange lines/dots displayed is the average probability acquired by training data set for each model and the blue lines/dots represents toxicity profile of the input compound depicting the probabilities of the input compound for the respective models. From the plot, we see that the SER alkaloid (Figure 3B) has a strong prediction confidence in the case of immunotoxicity as compared to that of the average probability model. The toxicological end points and pathways (models) were analyzed and are reported in Table S7. The data presented shows active prediction for carcinogenicity (0.55) and immunotoxicity (0.75) in case of AJM drug while for SER, the active predicted models were carcinogenicity (0.52), immunotoxicity (0.97) and mutagenicity (0.51) respectively. The immunotoxicity for the case of SER shows the highest probability score as compared to the other active models by both the drugs. Therefore, special attention should be given to these drugs for the predicted toxicity. However, for the remaining models like hepatotoxicity, cytotoxicity, Tox21 signaling and stressed pathways the predicted results shows inactive state as represented in the table. These results further led us to discovery the structural properties and parameters of the drugs.

**Figure 3.**
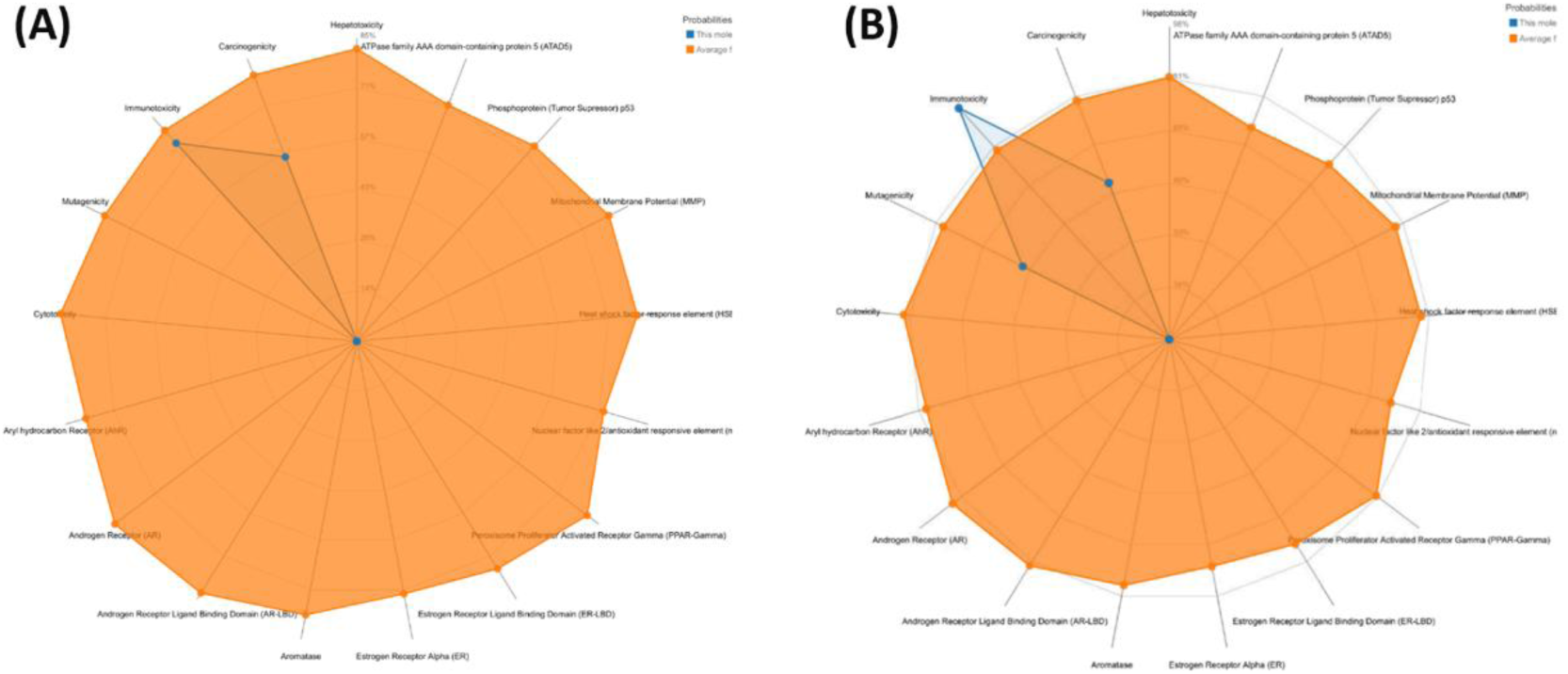
ProTox-II radar plot for (A) AJM and (B) SER drugs, illustrating the confidence of positive toxicity results compared to the average of its class.

### 3.3. Density Functional Theory (DFT)

DFT is a useful computational quantum mechanical tool for modeling electronic structure, electronic transition etc., for different molecules. The HOMO-LUMO energy gap (Figure 4, A and B) denotes the molecule stability index, chemical activity and bioactive property presented in Table 1. The HOMO energy is associated with the electron-donating ability required to remove an electron from the molecules ground state and is related with the ionization potential of the molecule whereas that of LUMO energy is related to the electron affinity and its ability to accept electrons. Therefore, the higher value of HOMO–LUMO energy gap for AJM results in high chemical stability and lower HOMO-LUMO gap for SER is associated with high reactivity.[54–56]

**Figure 4.**
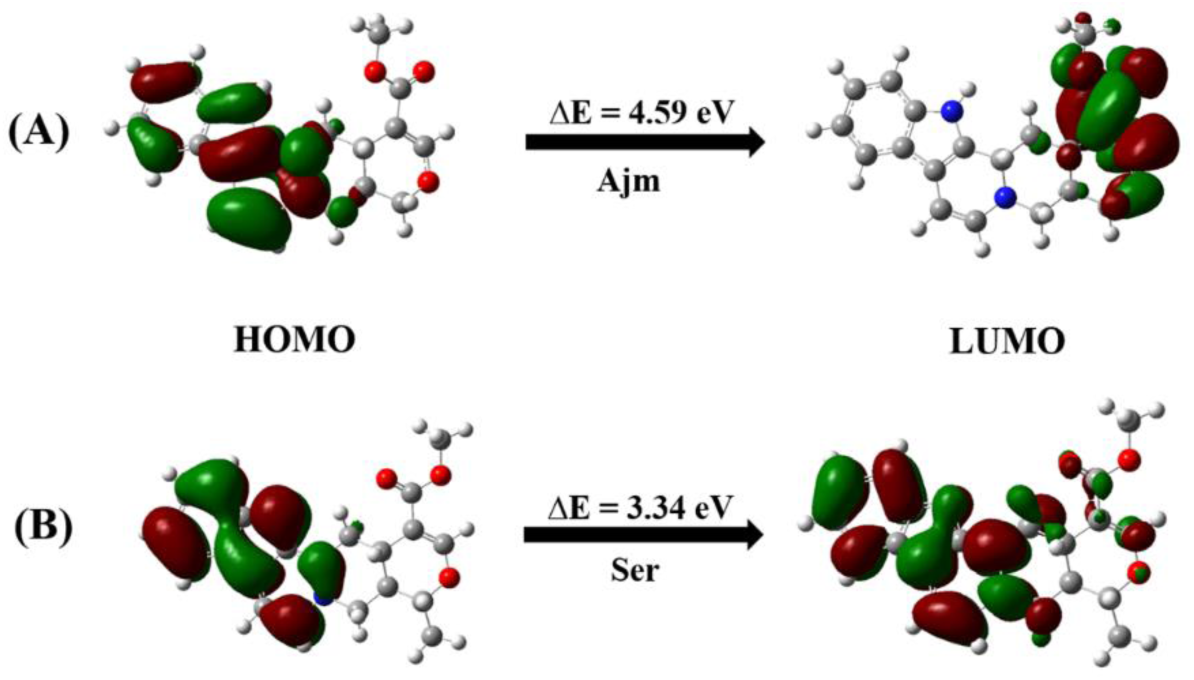
DFT plot of the highest occupied molecular orbital (HOMO) and the lowest unoccupied molecular orbital (LUMO) of (A) AJM and (B) SER.

**Table 1.**
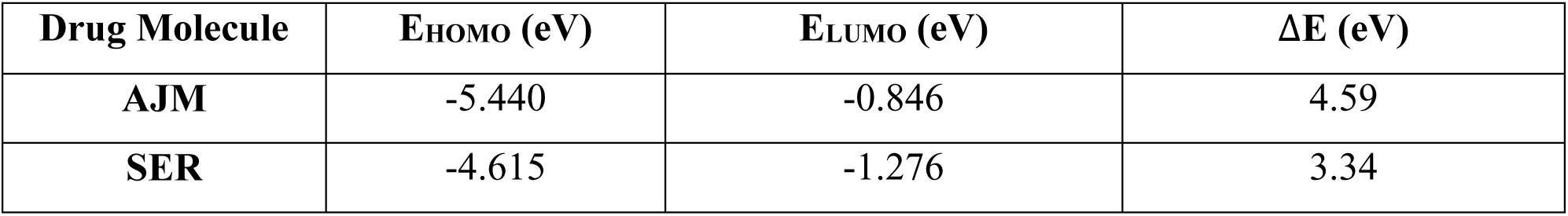
Parameters of AJM and SER, HOMO-LUMO energies and energy gap obtained from the optimized geometries.

The quantum chemical parameters of DFT descriptors were calculated from Koopmans’s theorem.[57–59] The theorem’s mathematical definitions were used to identify the drug reactivity (Table 2) such as ionization potential (*I*) and electronic affinity (*A*) as mentioned above. The chemical hardness (*η*) and softness (*ζ*) of the drug molecules suggest high and low resistance towards deformation in the electronic distribution during a reaction respectively.[56, 60, 61] The results obtained showed higher chemical hardness values and lower value of chemical softness for both the drug molecules. The electronegativity of the molecules indicate its ability to attract an electron which shows AJM having higher values as compared to that of SER. The chemical potential of the drugs were also determined which denotes energy that can be absorbed or released in a chemical reaction. The higher value of AJM (−3.14) reveals property of better electron donor ability over SER (−2.94) while the lesser negative value of SER shows good electron acceptor characteristic. Electrophilicity index of a molecule is a measure of an electrophile to accept an electron and its ability to resist exchange electron density with the surrounding in a reaction.[62, 63] The electrophilicity of a molecule can be classified as weak, moderate and strong depending on the value of its electrophilicity index.[64] Electrophiles having less than 0.8 eV are considered as weak, values between 0.8 to 1.5 eV are regarded moderate and above 1.5 eV are indicators of strong electrophiles. The values obtained from the investigated compounds signify strong and good electrophiles for both the cases, AJM having higher value as compared to that of SER. The values denote potent antimicrobial and anticancer properties of the drug molecules.[65] Similarly, nucleophiles can be regarded as strong (above 3.0 eV), moderate (between 2.0 to 3.0 eV) and weak (below 2.0 eV). The calculated drug molecules indicate weak nucleophilic property showing values below 2.0 eV. The maximum charge transfer was further calculated to determine the maximum propensity of the molecule to acquire electronic charge from the environment.[66] The result showed higher maximum charge transfer ability for SER as compared to that of AJM.

**Table 2.**
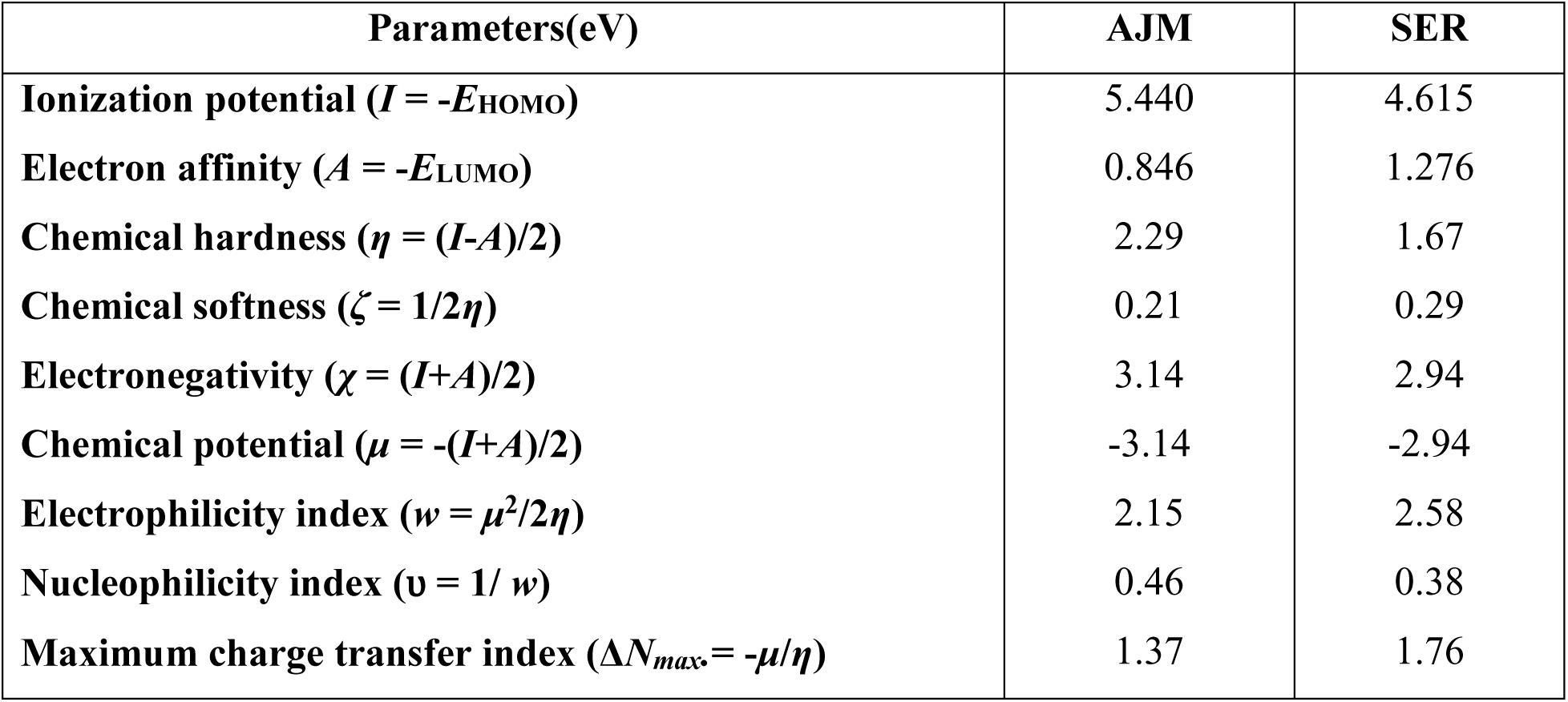
Quantum-chemical parameters of AJM and SER, computed with DFT at B3LYP/6-31G level.

| Parameters(eV) | AJM | SER |
| --- | --- | --- |
| **Ionization potential ($I = -E_{\text{HOMO}}$)** | 5.440 | 4.615 |
| **Electron affinity ($A = -E_{\text{LUMO}}$)** | 0.846 | 1.276 |
| **Chemical hardness ($\eta = (I-A)/2$)** | 2.29 | 1.67 |
| **Chemical softness ($\zeta = 1/2\eta$)** | 0.21 | 0.29 |
| **Electronegativity ($\chi = (I+A)/2$)** | 3.14 | 2.94 |
| **Chemical potential ($\mu = -(I+A)/2$)** | -3.14 | -2.94 |
| **Electrophilicity index ($w = \mu^2/2\eta$)** | 2.15 | 2.58 |
| **Nucleophilicity index ($v = 1/w$)** | 0.46 | 0.38 |
| **Maximum charge transfer index ($\Delta N_{\text{max}} = -\mu/\eta$)** | 1.37 | 1.76 |

### 3.4. Molecular Electrostatic Potential (MEP)

The optimized structures of the AJM and SER molecules were then used to determine the molecular electrostatic potential (MEP) to predict the chemical reactivity of the molecules toward positively and negatively charged reactants in their environment.[67] The diagram (Figure 5) provides a visual method for determining the relative polarity of the molecule, which determines a suitable position for nucleophilic and electrophilic attack. The color sequence of the electrostatic potential of the molecule at the surface increases in the following order, red < orange < yellow < green < blue.[68, 69] Figure 5A shows the negative potential phase as a dark red region located at the C=O bond, which is considered or preferred by the reactants as an electrophilic attack site. The positive potential site with blue color, corresponding to the nucleophilic attack site, is seen at the nitrogen atom of the pyrrole ring structure. Similarly, SER in Figure 5B shows a moderately positive charge site around the C=O bond of the structure and a stronger orange colored environment in the nitrogen atom of the pyrrole ring. The above discussion suggests the possibility of formation of bonds such as hydrogen bonds responsible for binding with the reactants around the electrophilic and nucleophilic attack sites.

**Figure 5.**
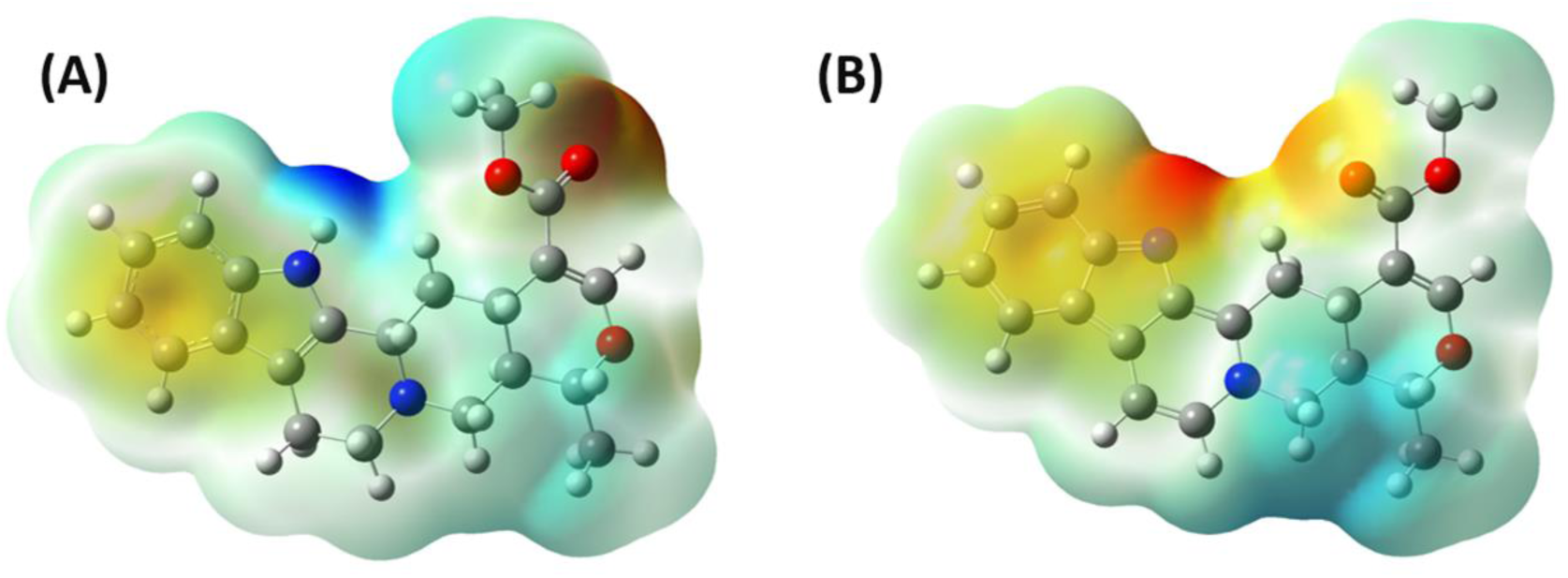
Molecular electrostatic potential plot of (A) AJM and (B) SER drug molecules.

The above theoretical data observed has assisted in understanding the molecular structure of the alkaloid drugs and its possibility of binding with bio-macromolecules like DNA.

### 3.4. Molecular docking study

Molecular docking study is an essential flexible docking method to predict the favorable conformations of the drug-DNA complex[21] and determine the structural configuration and binding modes of the drug to the target bio-macromolecules (proteins, DNA, etc.) with reasonable accuracy.[70, 71] The molecular docking study was initiated to reveal the binding site and bases involved in the binding between the alkaloids, AJM and SER with different forms of DNA, namely duplex, hairpin, triplex and quadruplex forms. Duplex DNA is two antiparallel aligned strands capable of encoding genetic information.[72] Figure 6 and 7 (A and B) show the interaction pattern of AJM and SER drug complexes with duplex DNA at the binding site of the minor groove for both cases. Figure 6 and 7 (C) shows the hydrogen bonding environment (surface) of AJM and SER duplex DNA complex showing the donor (purple) and acceptor (green) regions of the complex. Figure 6D shows how AJM forms conventional hydrogen bonds with dg14 (14th position of guanine) and dc7, carbon-hydrogen bonds with dg5, pi-pi T-shaped bonds with dg4, and pi-alkyl bonds with dc6. Figure 7D shows the conventional hydrogen bond SER with dg14 and dg15, the carbon-hydrogen bond with da8, the unfavorable acceptor-acceptor bond with dc16, and the pi-alkyl bond with dc7. The estimated free energy of binding was higher for SER duplex DNA complex than for AJM duplex DNA complex (Table 3), indicating that SER is a better binder for DNA. For all other cases such as hairpin duplex, triplex and quadruplex DNAs, the same pattern was seen in terms of binding free energy, i.e. SER showed higher values than AJM and thus better binding with the bio-macromolecule. The result is in agreement with that of DFT analysis: SER with a lower energy gap than AJM is more reactive and therefore binds better to the target (DNA). For the case of hairpin-alkaloid complexes (Figure 8 and 9), both the drug binds at the minor groove of the DNA with SER forming only one convention hydrogen bond with dt26 whereas AJM shows none. Triplex-drug complexes shows similar minor groove binding pattern as that of the previous complexes. Both the drug shows one conventional hydrogen bond each, dc5 for AJM (Figure 10) and dg12 for SER (Figure 11). For the case of drug-quadruplex complexes, the molecule was observed to bind and interact at the groove region of the G-quadruplex DNA (Figure 12 and 13). Both the molecules (AJM, SER) were found wrap around the top of the G-quadruplex DNA, AJM forming carbon-hydrogen bond with dg9, dg10 and dg14, pi-alkyl bond with dt12 and da13, pi-anion bond with dg15 whereas for SER, the interaction was through conventional hydrogen bond with dg8 and dg9, pi-anion bond with dg15, carbon hydrogen bond dg14 and pi-alkyl bond with da13. Table 3 also represents other various parameters involved in the binding between the bio-macromolecule and the drugs. AJM-triplex DNA, AJM-quadruplex DNA and SER-quadruplex DNA complexes have an inhibitory constant (Ki) greater than 100 μM, indicating that they are potent inhibitors.[73] The lower the Ki value of the drug, stronger is its binding affinity to the receptor. The docked complexes were analyzed in terms of hydrogen bonding, non-covalent bonding and hydrophobic interaction between the drugs and the various DNAs. The obtained results show that the drugs successfully bind to DNA and are potential DNA binders. This docking information can help researchers to perform further experimental analysis of alkaloid drugs with DNA.

**Figure 6.**
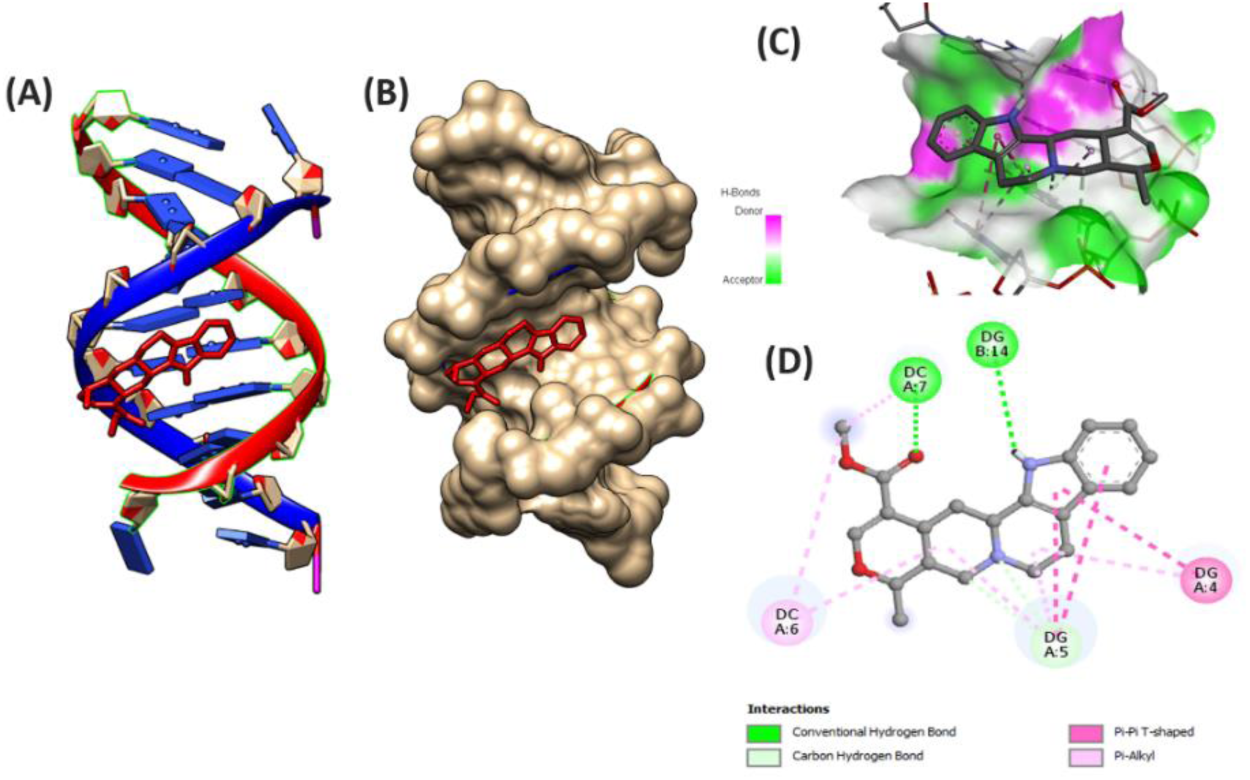
Molecular docking picture representation of AJM-duplex DNA complex (A) ribbon (DNA) and stick (AJM) form (B) hydrophobic (DNA) and stick (AJM) form (C) ligand in hydrogen vicinity of the bio-macromolecule and (D) 2D representation of the ligand-DNA interaction.

**Figure 7.**
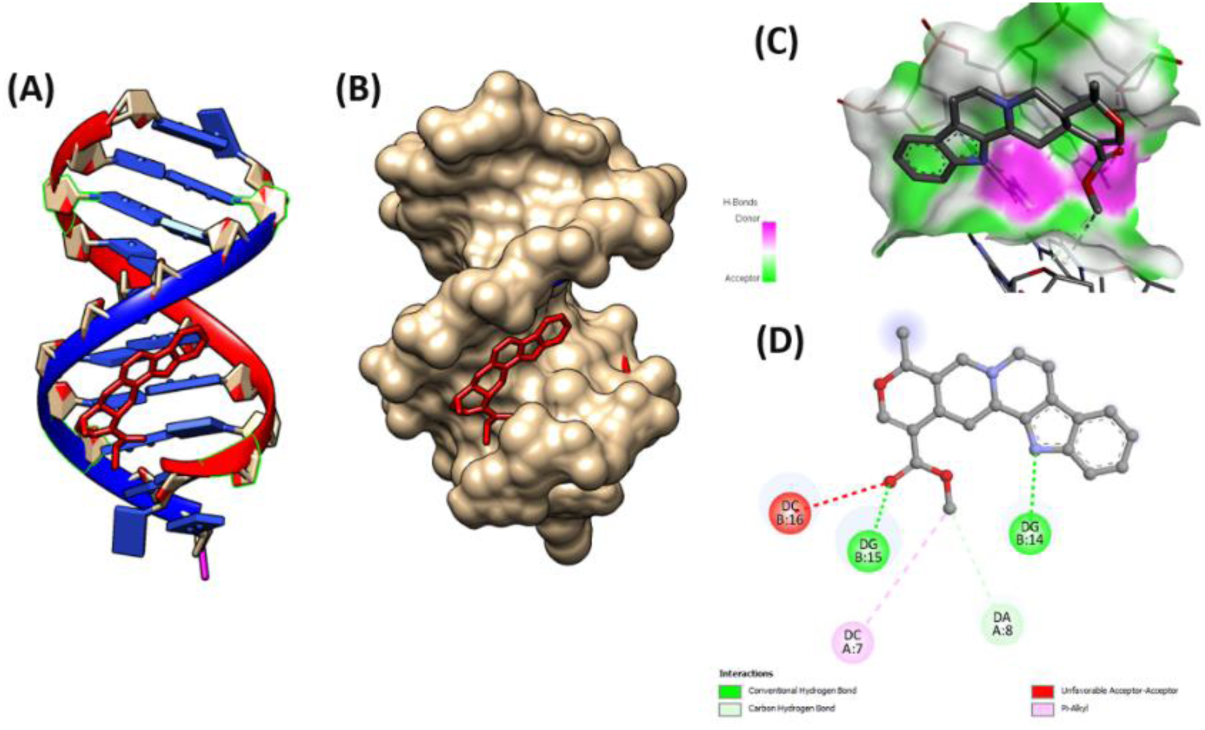
Molecular docking picture representation of SER-duplex DNA complex (A) ribbon (DNA) and stick (SER) form (B) hydrophobic (DNA) and stick (SER) form (C) ligand in hydrogen vicinity of the bio-macromolecule and (D) 2D representation of the ligand-DNA interaction.

**Figure 8.**
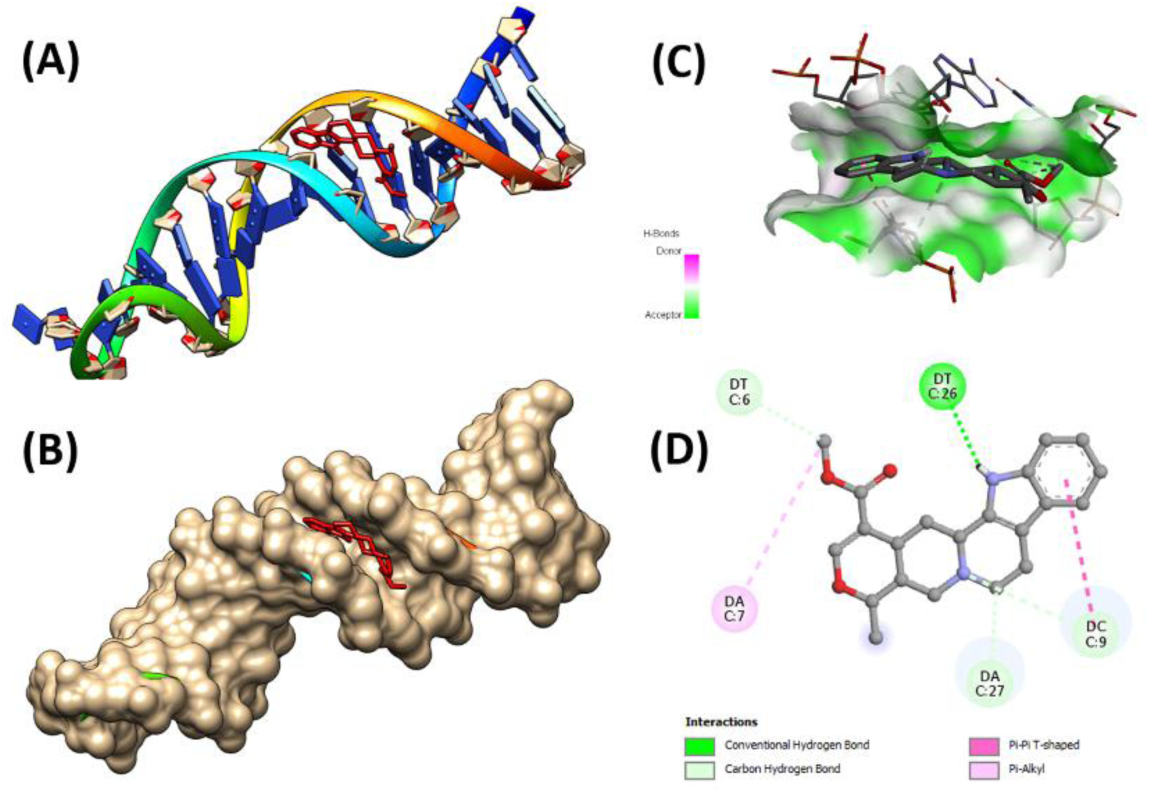
Molecular docking picture representation of AJM-hairpin DNA complex. (A) ribbon (DNA) and stick (AJM) form (B) hydrophobic (DNA) and stick (AJM) form (C) ligand in hydrogen vicinity of the bio-macromolecule and (D) 2D representation of the ligand-DNA interaction.

**Figure 9.**
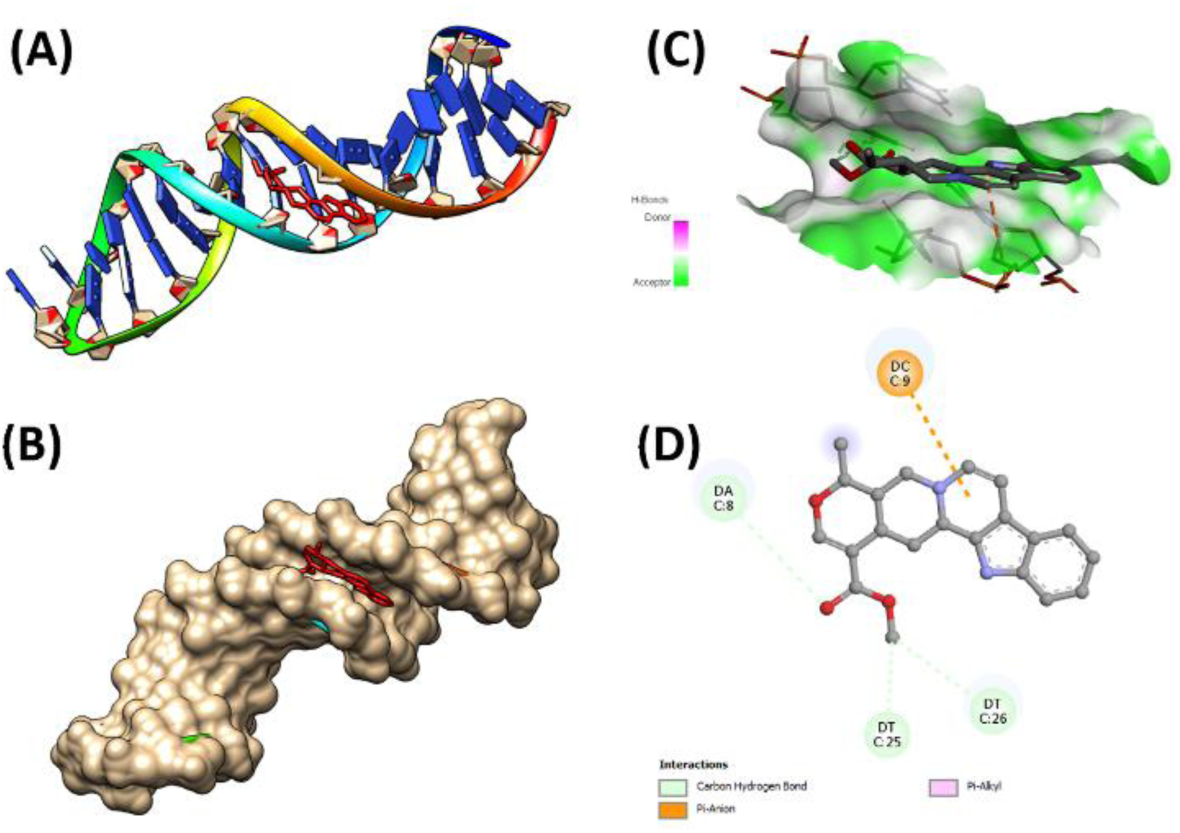
Molecular docking picture representation of SER-hairpin DNA complex. (A) ribbon (DNA) and stick (SER) form (B) hydrophobic (DNA) and stick (SER) form (C) ligand in hydrogen vicinity of the bio-macromolecule and (D) 2D representation of the ligand-DNA interaction.

**Figure 10.**
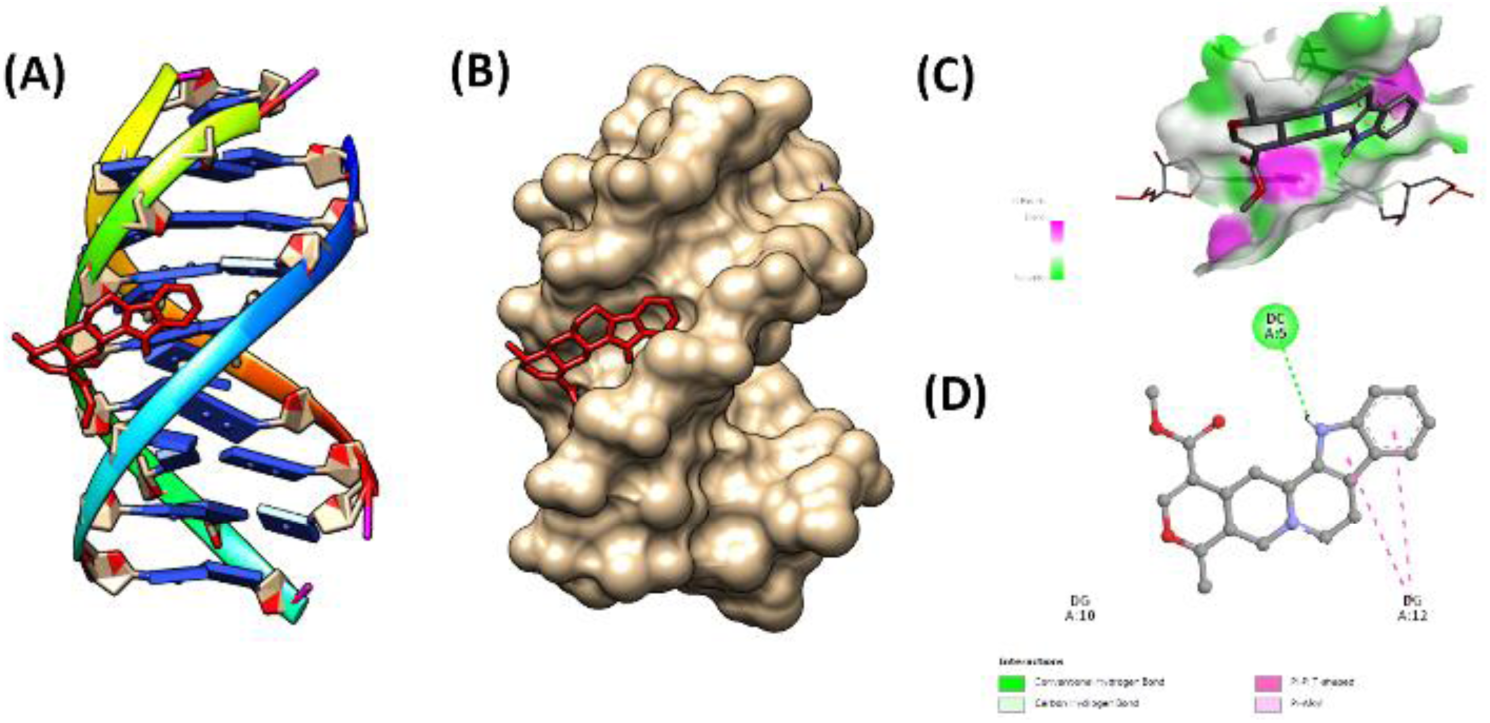
Molecular docking picture representation of AJM-triplex DNA complex. (A) ribbon (DNA) and stick (AJM) form (B) hydrophobic (DNA) and stick (AJM) form (C) ligand in hydrogen vicinity of the bio-macromolecule and (D) 2D representation of the ligand-DNA interaction.

**Figure 11.**
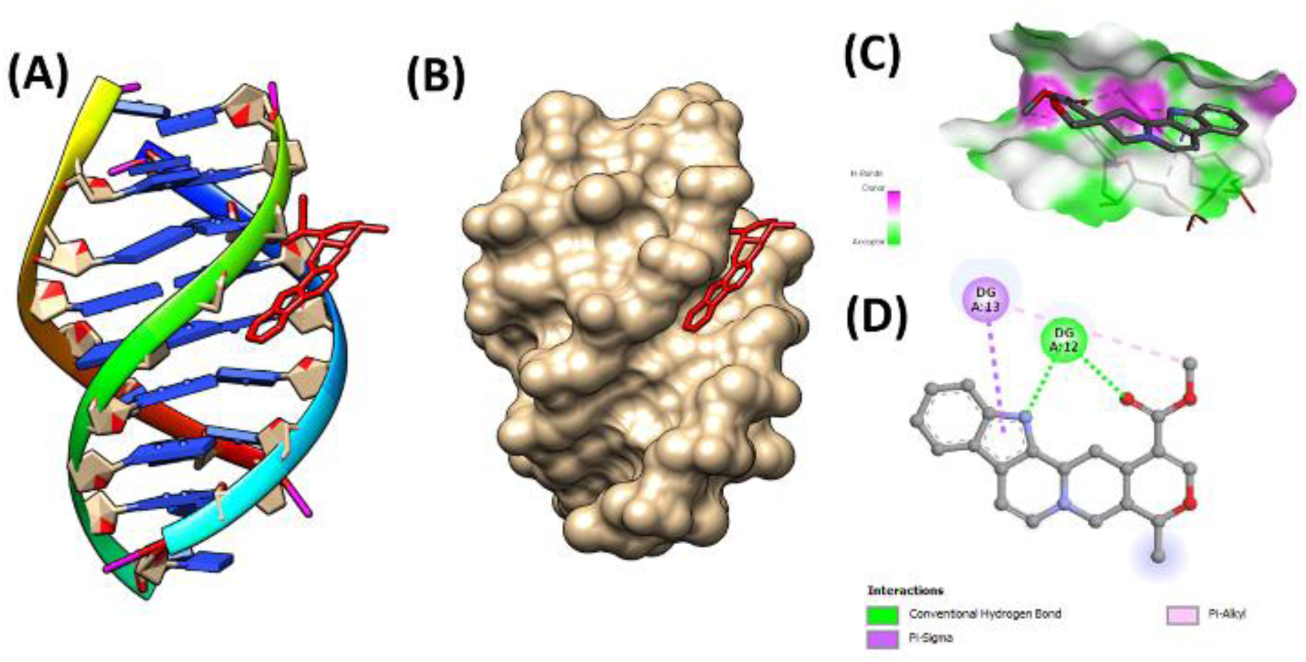
Molecular docking picture representation of SER-triplex DNA complex. (A) ribbon (DNA) and stick (SER) form (B) hydrophobic (DNA) and stick (SER) form (C) ligand in hydrogen vicinity of the bio-macromolecule and (D) 2D representation of the ligand-DNA interaction.

**Figure 12.**
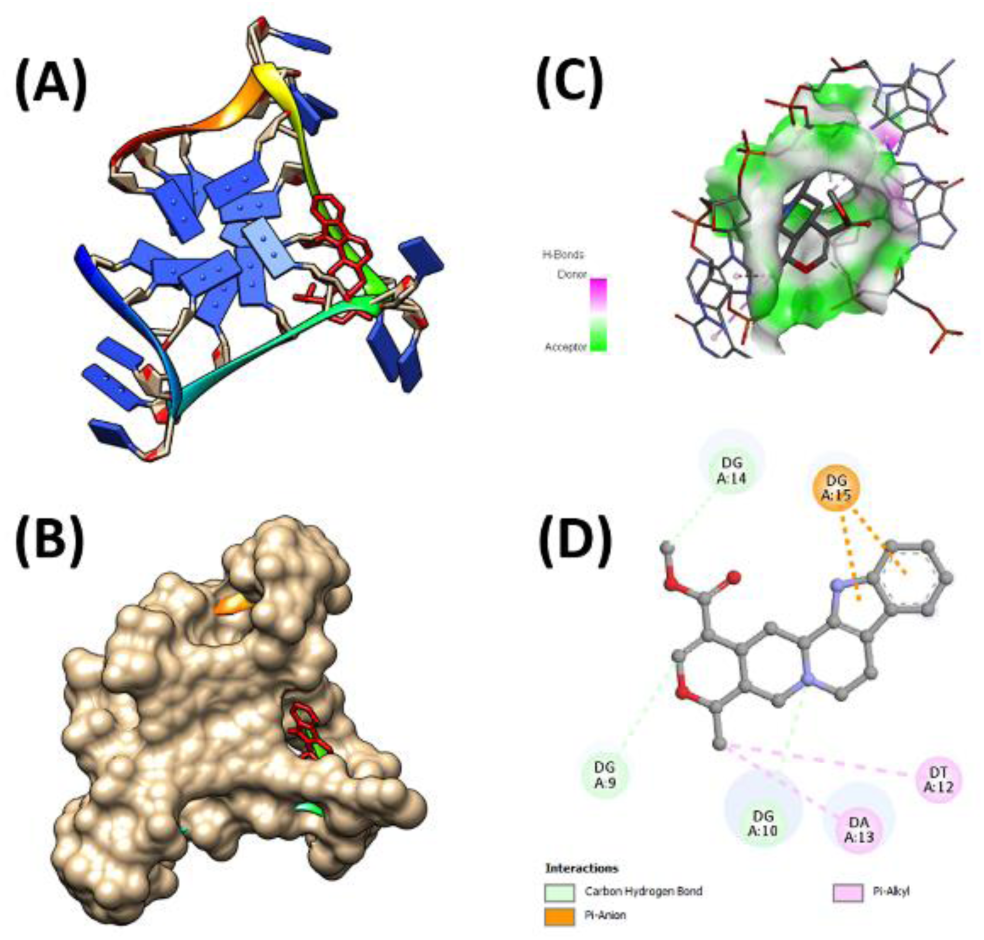
Molecular docking picture representation of AJM-quadruplex DNA complex. (A) ribbon (DNA) and stick (AJM) form (B) hydrophobic (DNA) and stick (AJM) form (C) ligand in hydrogen vicinity of the bio-macromolecule and (D) 2D representation of the ligand-DNA interaction.

**Figure 13.**
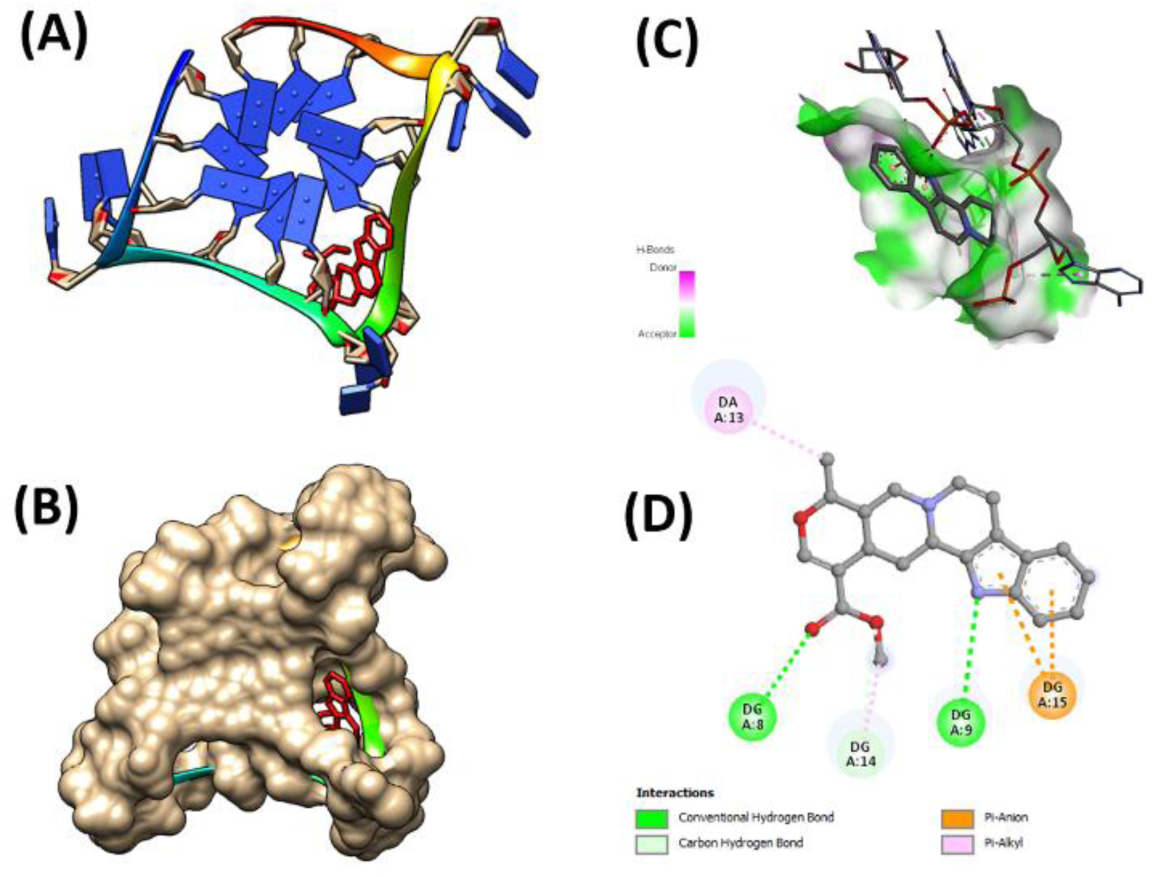
Molecular docking picture representation of SER-quadruplex DNA complex. (A) ribbon (DNA) and stick (SER) form (B) hydrophobic (DNA) and stick (SER) form (C) ligand in hydrogen vicinity of the bio-macromolecule and (D) 2D representation of the ligand-DNA interaction.

**Table 3.**
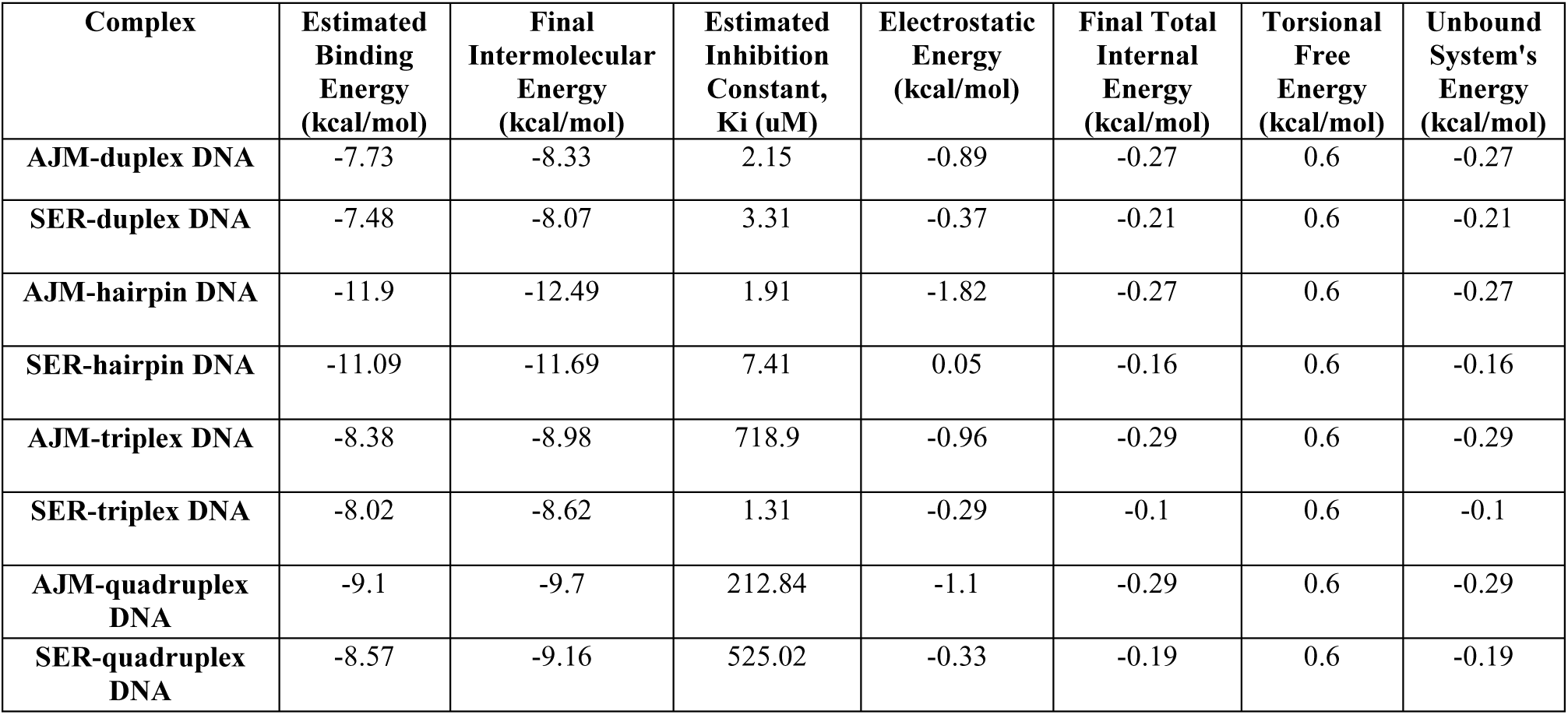
Autodock results on complexes of AJM and SER with different forms of DNA.

## 4. Conclusions

In this paper, pilot screening of indole alkaloid molecules AJM and SER (commonly found in medicinal plant of Nagaland) is studied using SwissADME, potential toxicity (ProTox-II), Gaussian studies and binding studies with different forms of DNA using molecular docking studies. The drug properties (AJM, SER) obtained from ADME analysis show that they fall within the acceptable range/parameters of a standard drug. A large number of in silico toxicological endpoints were predicted showing active prediction for carcinogenicity and immune-toxicity for AJM & carcinogenicity, immune-toxicity and mutagenicity for SER. The DFT study provides the optimized drug details that include the structure of alkaloids with their energies, quantum chemical parameters, and molecular electrostatic potential to predict the chemical reactivity of the molecules. Docking information with the different DNA models indicates successful binding between the ligand and the bio-macromolecules. This pre-modeled theoretical analysis can be considered as a powerful computational result to reduce the workload and cost required for the development of effective medicinal compounds, which are helpful for a better understanding of their bioactive interaction mechanisms with different forms of DNA.

## Ethical Approval

The study requires no ethical approval. The manuscript is in compliance with ethical standards.

## Consent to Participate

NA

## Consent to Publish

Consent to publish has been received from all authors.

## Authors Contributions

Performing experiments; Data analyses, Figures and Tables; Manuscript writing: Aben Ovung Partial manuscript writing and all associated files preparation: A. Mavani Gaussian analyses: Gourisankar Roymahapatra Concept, overall guidance, project management and research fund acquisition: Jhimli Bhattacharyya

## Availability of data and materials

Research data will be made available from the corresponding author, upon reasonable request.

## Supplementary Information

**Table S1:** Physicochemical Properties of indole alkaloids, AJM and SER.

| Small molecule | Num. heavy atoms | Num. arom. heavy atoms | Fraction Csp3 | Num. rotatable bonds | Num. H-bond acceptors | Num. H-bond donors | Molar refractivity | TPSA (Å<sup>2</sup>) |
| --- | --- | --- | --- | --- | --- | --- | --- | --- |
| AJM | 26 | 9 | 0.48 | 2 | 4 | 1 | 103.47 | 54.56 |
| SER | 26 | 13 | 0.33 | 2 | 4 | 0 | 99.59 | 53.35 |

**Table S2:** Lipophilicity characteristics of indole alkaloids, AJM and SER.

| Small molecule | iLOGP | XLOGP3 | WLOGP | MLOGP | SILICOS-IT | Consensus Log *P*<sub>o/w</sub> |
| --- | --- | --- | --- | --- | --- | --- |
| AJM | 3.20 | 2.75 | 2.47 | 2.13 | 2.78 | 2.67 |
| SER | 3.33 | 2.58 | 3.45 | 2.21 | 2.78 | 2.87 |

**Table S3:** Water solubility characteristics of indole alkaloids, AJM and SER.

| Small molecule | ESOL |  |  |  | Ali |  |  |  | SILICOS- IT |  |  |  |
| --- | --- | --- | --- | --- | --- | --- | --- | --- | --- | --- | --- | --- |
|  | Log S | Solubility |  | Class | Log S | Solubility |  | Class | Log S | Solubility |  | Class |
|  |  | mg/ml | mol/L |  |  | mg/ml | mol/L |  |  | mg/ml | mol/L |  |
| AJM | -3.88 | 4.63e-02 | 1.31e-04 | S | -3.55 | 9.92e-02 | 2.81e-04 | S | -4.38 | 1.47e-02 | 4.06e-05 | MS |
| SER | -3.86 | 4.77e-02 | 1.37e-04 | S | -3.35 | 1.56e-01 | 4.48e-04 | S | -4.92 | 4.14e-03 | 1.19e-05 | MS |

**Table S4:** Pharmacokinetics parameters of indole alkaloids, AJM and SER.

| Small molecule | GI absorption | BBB permeant | P-gp substrate | CYP1A2 inhibitor | CYP2C19 inhibitor | CYP2C9 inhibitor | CYP2D6 inhibitor | CYP3A4 inhibitor | Log *K*<sub>p</sub> (Skin Permeation) (cm/s) |
| --- | --- | --- | --- | --- | --- | --- | --- | --- | --- |
| AJM | High | Yes | Yes | No | No | No | Yes | No | -6.50 |
| SER | High | Yes | No | No | Yes | No | Yes | Yes | -6.59 |

**Table S5:**
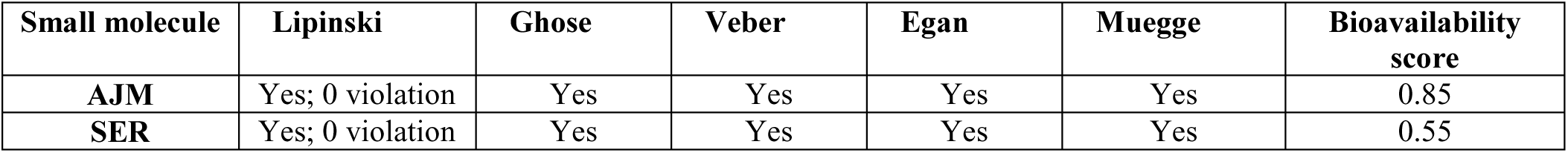
Druglikeness rule and Bioavailability score of indole alkaloids, AJM and SER.

| Small molecule | Lipinski | Ghose | Veber | Egan | Muegge | Bioavailability score |
| --- | --- | --- | --- | --- | --- | --- |
| AJM | Yes; 0 violation | Yes | Yes | Yes | Yes | 0.85 |
| SER | Yes; 0 violation | Yes | Yes | Yes | Yes | 0.55 |

**Table S6:** Medicinal Chemistry properties of indole alkaloids, AJM and SER.

| Small molecule | Pains | Brenk | Leadlikeness | Synthetic accessibility |
| --- | --- | --- | --- | --- |
| AJM | 1 alert: indol_3yl_alk | 0 | No; 1 violation: MW>350 | 4.57 |
| SER | 0 | 0 | Yes | 4.32 |

**Table S7:**
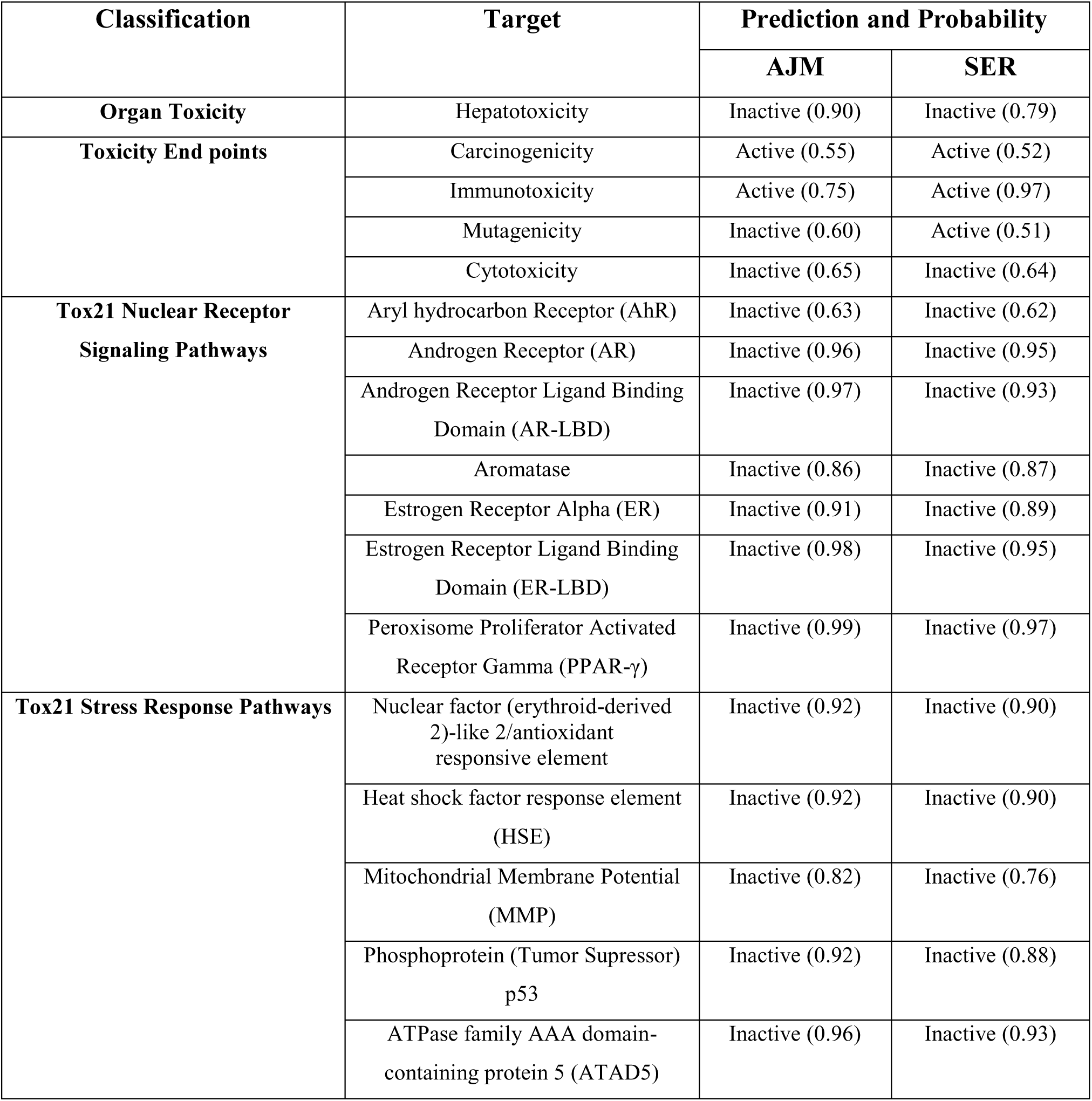
Toxicity Model reports for the alkaloids, AJM and SER.

| Classification | Target | Prediction and Probability |  |
| --- | --- | --- | --- |
|  |  | AJM | SER |
| Organ Toxicity | Hepatotoxicity | Inactive (0.90) | Inactive (0.79) |
| Toxicity End points | Carcinogenicity | Active (0.55) | Active (0.52) |
|  | Immunotoxicity | Active (0.75) | Active (0.97) |
|  | Mutagenicity | Inactive (0.60) | Active (0.51) |
|  | Cytotoxicity | Inactive (0.65) | Inactive (0.64) |
| Tox21 Nuclear Receptor Signaling Pathways | Aryl hydrocarbon Receptor (AhR) | Inactive (0.63) | Inactive (0.62) |
|  | Androgen Receptor (AR) | Inactive (0.96) | Inactive (0.95) |
|  | Androgen Receptor Ligand Binding Domain (AR-LBD) | Inactive (0.97) | Inactive (0.93) |
|  | Aromatase | Inactive (0.86) | Inactive (0.87) |
|  | Estrogen Receptor Alpha (ER) | Inactive (0.91) | Inactive (0.89) |
|  | Estrogen Receptor Ligand Binding Domain (ER-LBD) | Inactive (0.98) | Inactive (0.95) |
| | Peroxisome Proliferator Activated Receptor Gamma (PPAR-γ) | Inactive (0.99) | Inactive (0.97) |
| Tox21 Stress Response Pathways | Nuclear factor (erythroid-derived 2)-like 2/antioxidant responsive element | Inactive (0.92) | Inactive (0.90) |
|  | Heat shock factor response element (HSE) | Inactive (0.92) | Inactive (0.90) |
|  | Mitochondrial Membrane Potential (MMP) | Inactive (0.82) | Inactive (0.76) |
|  | Phosphoprotein (Tumor Suppressor) p53 | Inactive (0.92) | Inactive (0.88) |
|  | ATPase family AAA domain-containing protein 5 (ATAD5) | Inactive (0.96) | Inactive (0.93) |

